# Synergistic therapeutic effects of endolysin BNT331 and a *Lactobacillus crispatus* consortium in a human vagina chip model of bacterial vaginosis

**DOI:** 10.1101/2025.11.05.686849

**Authors:** Aakanksha Gulati, Alicia C. Jorgenson, Ashleigh Williams, Hassan Rhbiny, Lenka Podpera Tisakova, Lorenzo Corsini, Mahya Fazel-Zarandi, Jiawu Xu, Joseph Elsherbini, Ola Gutzeit, Anna Stejskalova, Nina Teresa LoGrande, Briah Cooley Demidkina, Yuncheng Man, Girija Goyal, Douglas S. Kwon, Caroline M. Mitchell, Abidemi Junaid, Donald E. Ingber

## Abstract

Bacterial vaginosis (BV) is a vaginal infection caused by an imbalance in the vaginal microbiome characterized by a decrease in healthy bacteria dominated by *Lactobacillus crispatus* along with a concomitant increase in dysbiotic bacteria, such as *Gardnerella*. Current treatments commonly fail to fully eradicate BV, and the lack of more effective therapies is due in part to the absence of relevant human models. Here, we used a human vagina-on-a-chip (Vagina Chip) microfluidic culture device that has been previously shown to faithfully recapitulate the human vaginal microenvironment as well as the inflammatory and injurious effects of *G. vaginalis* to test the therapeutic efficacy of a consortium of *L. crispatus* alone or in combination with endolysin BNT331, which specifically targets and lyses *Gardnerella*. These studies revealed that when *L. crispatus* was added alone, it suppressed inflammation even though it failed to engraft or displace the *G. vaginalis* bacteria. In contrast, BNT331 effectively killed *Gardnerella* in dysbiotic Vagina Chip. Importantly, the combined administration of both treatments resulted in restoration of a healthier vaginal microenvironment, as indicated by higher engraftment of *L. crispatus* on-chip, inhibition of *G. vaginalis*, and a reduction in inflammation. Similar effects of this combined treatment were observed when administered to Vagina Chips infected with vaginal swab samples from BV patients. These data suggest that combination of a live biotherapeutic product composed of a *L. crispatus* consortium with a potent antimicrobial agent that targets *G. vaginalis*, such as BNT331, may offer an effective therapeutic strategy for patients with BV.

**One Sentence Summary:** Endolysin BNT331 combined with *Lactobacillus crispatus* reduces bacterial load and suppresses inflammation in a human vagina chip model of bacterial vaginosis.

## INTRODUCTION

Bacterial vaginosis (BV) is one of the most common vaginal diseases caused by an imbalance in the vaginal microbiome, which not only causes vaginal inflammation and a malodorous discharge, but also has been shown to pre-dispose women to an increased risk of pre-term birth and adverse pregnancy outcomes as well as sexually transmitted infections (STIs), infertility, and cervical cancer (*1–8*). BV is characterized by a decrease in *Lactobacillus crispatus* and other *Lactobacilli* that dominate the optimal (healthy) microbiome along with an increase in non-optimal (dysbiotic) consortia of *Gardnerella vaginalis* and other anaerobic bacterial species (e.g., *Prevotella sp.*, *Atopobium vaginae*, *Sneathia)* (*9, 10*). The current treatment for BV is antibiotics, administered either orally or intra-vaginally; however, they are often ineffective over time resulting in recurrent BV (*11*). Hence, there is a dire need to develop better treatment strategies to effectively treat BV and reduce the recurrence of this disease.

A healthy vaginal microbiome is dominated by *Lactobacillus* species, such as *L. crispatus*, *L. gasseri*, *L. iners*, *L. jensenii*, *L. reuteri* and *L. vaginalis* with *L. crispatus* being the most common one (*12*). *Lactobacillus* sp. produce lactic acid, induce the production of anti-microbial peptides, and stimulate the production of mucus by the cervico-vaginal tissue (*13–15*), thereby protecting the vagina from sporadic infections. Moreover, patients with BV exhibit a shift in the dominant consortia of the vaginal microbiome so that *Lactobacilli* become less prevalent and anaerobic species often containing *G. vaginalis* take over. The current standard of care involves administering antibiotics, such as metronidazole or clindamycin, either orally or intra-vaginally; however, high recurrence of BV is often observed after these treatments. Recently, attempts to treat recurrent BV with vaginal microbial transplantation (VMT) from a healthy donor to BV patients have been found to be successful, but VMT poses risks of transfer of other anti-microbial resistant microbes, undetected pathogens such as viruses, and they even pose a risk of unwanted pregnancy due to the presence of sperm (*16*). Thus, a new approach that is being explored to treat BV without producing these side effects is to use a live biotherapeutic product (LBP) composed of a consortia of *Lactobacillus* sp. to reverse the dysbiotic microbiome to a healthy *L. crispatus*- dominated state (*17*). However, others have pursued the possibility that BV could be treated by inhibiting the growth of potential pathogens, such as *G. vaginalis,* particularly using therapies that overcome bacterial resistance. The goal of both approaches is to develop a therapy that can not only reverse the BV phenotype but do so in a sustained manner.

In the present study, we leveraged a recently described human vagina-on-a-chip (Vagina Chip) microfluidic culture model of BV that is driven by engrafting *G. vaginalis* on-chip (*18*) to compare the therapeutic potential of a targeted endolysin (BNT331), which has been shown to specifically kill *G. vaginalis* in both suspension and in biofilms *in vitro* (*19*), and a human patient-derived *L. crispatus* consortium that was previously shown to engraft in human Vagina Chips. Our results show that BNT331 killed *G. vaginalis* in the Vagina Chip and inhibited its growth. Administration of *a L. crispatus* consortium alone suppressed inflammation, but it did not reduce the dysbiotic bacterial load. In contrast, combining both therapeutic approaches lead to reduction of *Gardnerella*, engraftment of *L. crispatus* and reduction of inflammation at the same time, indicative of a more healthy vaginal microbiome. Moreover, we demonstrate similar results when the chips are engrafted with a more clinically relevant, dysbiotic microbiome obtained from vaginal swab samples collected from BV patients.

## RESULTS

### Mimicking the healthy and dysbiotic vaginal microenvironment on-chip

The Vagina Chip is a microfluidic device with two microchannels separated by a porous membrane that contains a stratified human vaginal epithelium cultured on the top of the membrane and a stroma containing human vaginal stromal fibroblasts grown on the lower side of the same membrane (**Fig. 1A**). To mimic healthy (optimal) and dysbiotic (non-optimal) states of the human vagina in vitro, we inoculated the Vagina Chip with either *L. crispatus* or *G. vaginalis,* respectively, and both were able to successfully engraft on-chip when quantified by counting colony forming units (CFUs) after enzymatically degrading the epithelium (**Fig. 1A-C**). Immunofluorescence microscopic analysis confirmed that the vaginal epithelium exposed to *L. crispatus* remained healthy as indicated by retention of a basal layer of CK-5 expressing cells and the presence of intact ZO-1-containing tight junctions (ZO-1), whereas exposure to *G. vaginalis* resulted in a major decrease in CK-5 basal cells and complete loss of tight junctions when analyzed 72 h after inoculation (**Fig. 1C**). Importantly, *G. vaginalis*, but not *L. crispatus,* also induced significant cytotoxicity as measured using an LDH assay (**Fig. 1D**) as well as increased secretion of multiple pro-inflammatory cytokines (IL-6, IL- 8, IL-1a, IL-1b, TNF-a) compared to uninfected vaginal epithelial cells on-chip (**Fig. 1E**). These results align with the clinical finding that BV induces damage and inflammation in the vagina, whereas the presence of *L. crispatus* is associated with a healthy non-inflammatory vaginal environment. Altogether, these data show that the Vagina Chip can recapitulate both healthy and dysbiotic vaginal environments *in vitro*.

**Fig. 1.**
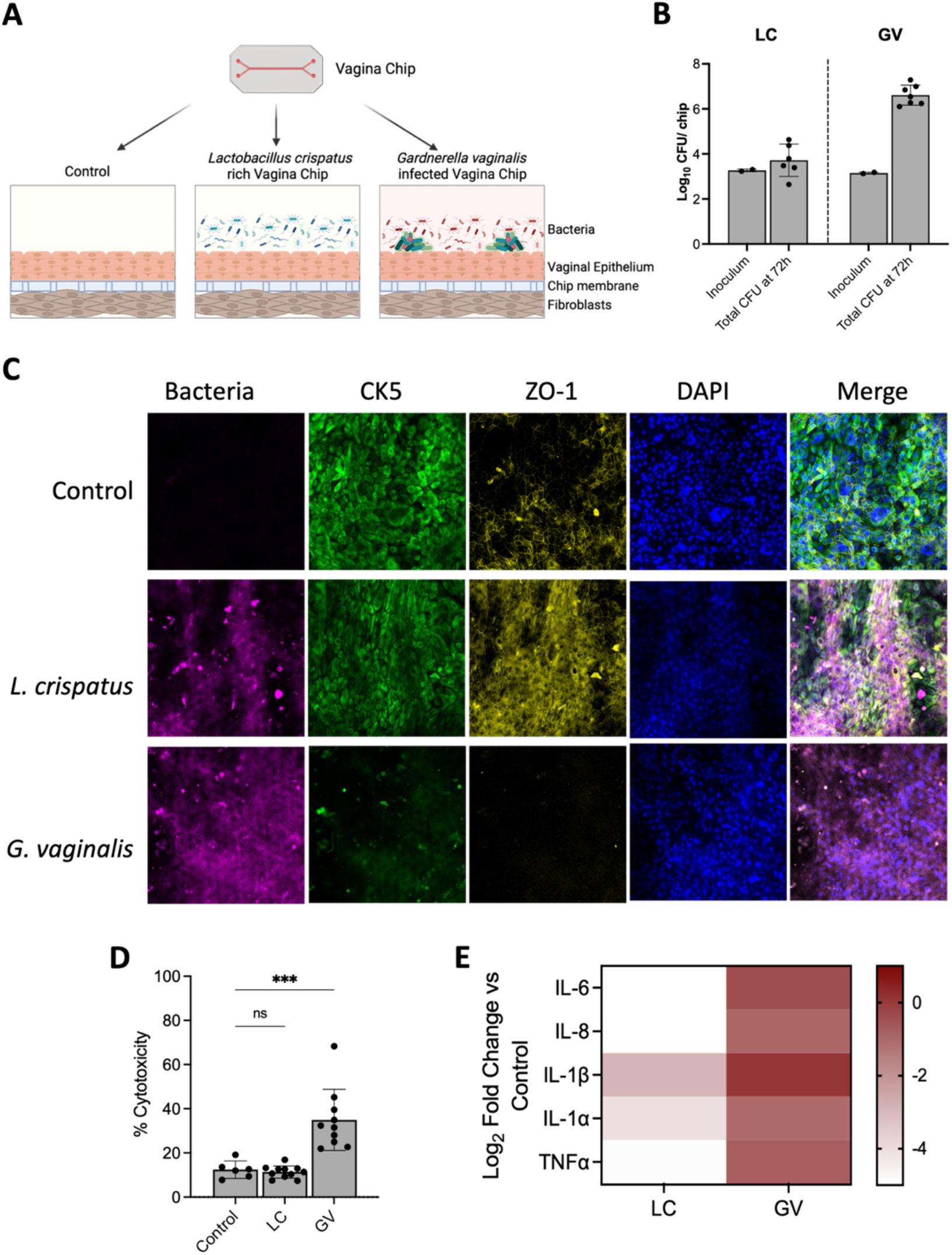
The healthy and dysbiotic bacteria engraft on the Vagina Chip; dysbiosis induces pro-inflammation and cell death in the Vagina Chip. A) Schematic representation of a Vagina Chip inoculated with *Lactobacillus crispatus* or *Gardnerella vaginalis.* **B)** Total bacterial load was calculated in Vagina Chips inoculated with *Lactobacillus crisp*atus (LC) or *Gardnerella vaginalis* (GV) at 72 h by enumerating bacterial CFU in the outflows and those adhered to the vaginal epithelial cells. Inoculum depicts the bacteria seeded in the Vagina Chip. Bar graph represents Mean ± SD from 2 independent experiments. **C)** Representative confocal images depicting the basal layer marker of vaginal epithelium, cytokeratin-5 (CK5) and the tight junction protein ZO-1 in uninfected (control) Vagina Chips or Vagina Chips inoculated with *L. crispatus* or *G. vaginalis* at 72 h. **D)** Cytotoxicity at 72h of inoculation of Vagina Chips with *L. crispatus* (LC) or G. vaginalis (GV) was measured using LDH assay. Bar graphs represent Mean ± SD from 3 independent experiments. P-value is calculated compared to uninfected control Vagina Chip one-way ANOVA. **E)** Pro-inflammatory cytokines were analyzed at 72h of inoculation of Vagina Chips with *L. crispatus* (LC) or *G. vaginalis* (GV) using Luminex-based assay. Log2 Fold change was calculated compared to the cytokine levels in uninoculated control Vagina Chip.

### *L. crispatus* reduces inflammation and cytotoxicity in *G. vaginalis*-infected Vagina Chips

*Lactobacillus* containing live biotherapeutic products (LBPs) are being explored as potential BV therapeutics because the healthy vaginal environment is dominated by *L. crispatus* bacteria, which are believed to protect against vaginal infections (*10*). We therefore explored whether a *L. crispatus* consortium containing multiple optimal strains can reverse the inflammation and cell injury induced by a *G. vaginalis-*containing dysbiotic microbiome in the human Vagina Chip (**Fig. 2A**). However, this treatment did not reduce the number of *G. vaginalis* bacteria (**Fig. 2B**), although it did suppress *G. vaginalis*-induced inflammation (**Fig 2C**) and this was associated with reduced cytotoxicity (**Fig. 2D**). However, to be able to fully restore a healthy vaginal microbiome and prevent the common recurrence of BV, it also will likely be necessary to lower the levels of *G. vaginalis*.

**Fig. 2.**
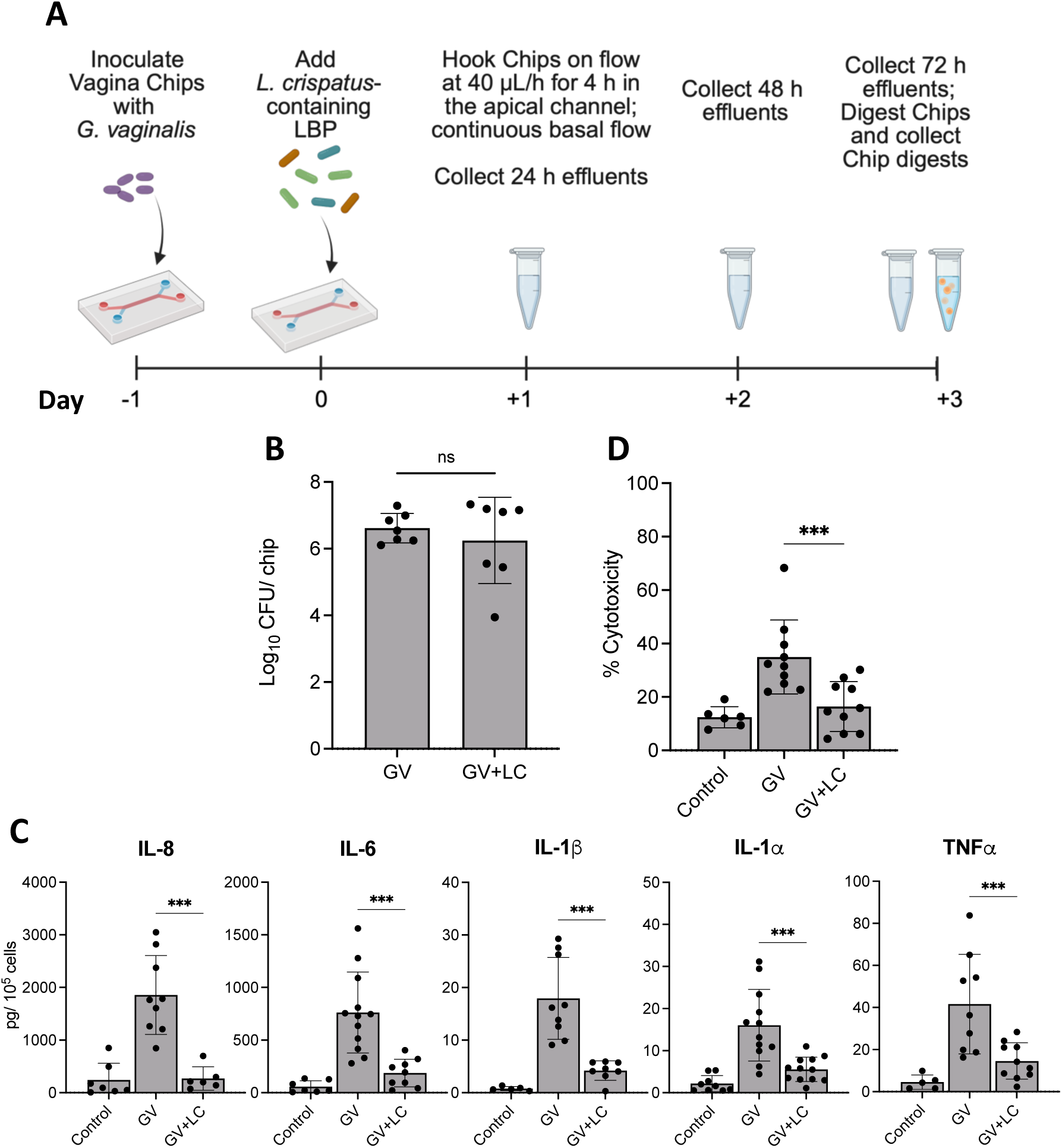
*L. crispatus* reduces *G. vaginalis*-induced inflammation but does not reduce *G. vaginalis* established in the Vagina Chip. A) Pictorial representation of the experimental workflow. *G. vaginalis*-infected Vagina Chips were treated with *L. crispatus* consortium (LC) 24 h after infection. CFU of G. vaginalis (GV) was enumerated 72 h after LC treatment. **C)** Pro-inflammatory cytokines in the outflows were detected after 72 h of LC treatment of GV-infected Vagina Chips using Luminex-based assay. **D)** Cytotoxicity was measured using LDH assay after 72 h of LC- treatment of GV-infected Vagina Chips. **(B-D)** Bar graphs represent Mean± SD for 3 independent experiments. P- values were calculated using one-way ANOVA.

### BNT331 abrogates G. vaginalis in the Vagina Chip

BNT331 is an engineered endolysin that is specifically designed to target *G. vaginalis* and induce its killing (*19*). When we incubated Vagina Chips with *G. vaginalis* for 24 hours and then treated them with 10, 25, 50 or 100 µg/mL of BNT331, we found that the 50 µg/mL dose completely abolished *G. vaginalis* from the Vagina Chip by 72 hours (**Fig. 3A**). As expected, this near total removal of *G. vaginalis* from the chip also resulted in a decrease in inflammation (**Fig. 3B**) and cytotoxicity (**Fig. 3C**). Importantly, similar results were obtained when we infected human Vagina Chips with complex dysbiotic microbiome isolated from vaginal swabs collected from BV patients, which has only a low level of *L. crispatus* (**fig. S2**) and then treated them with BNT331.

**Fig. 3.**
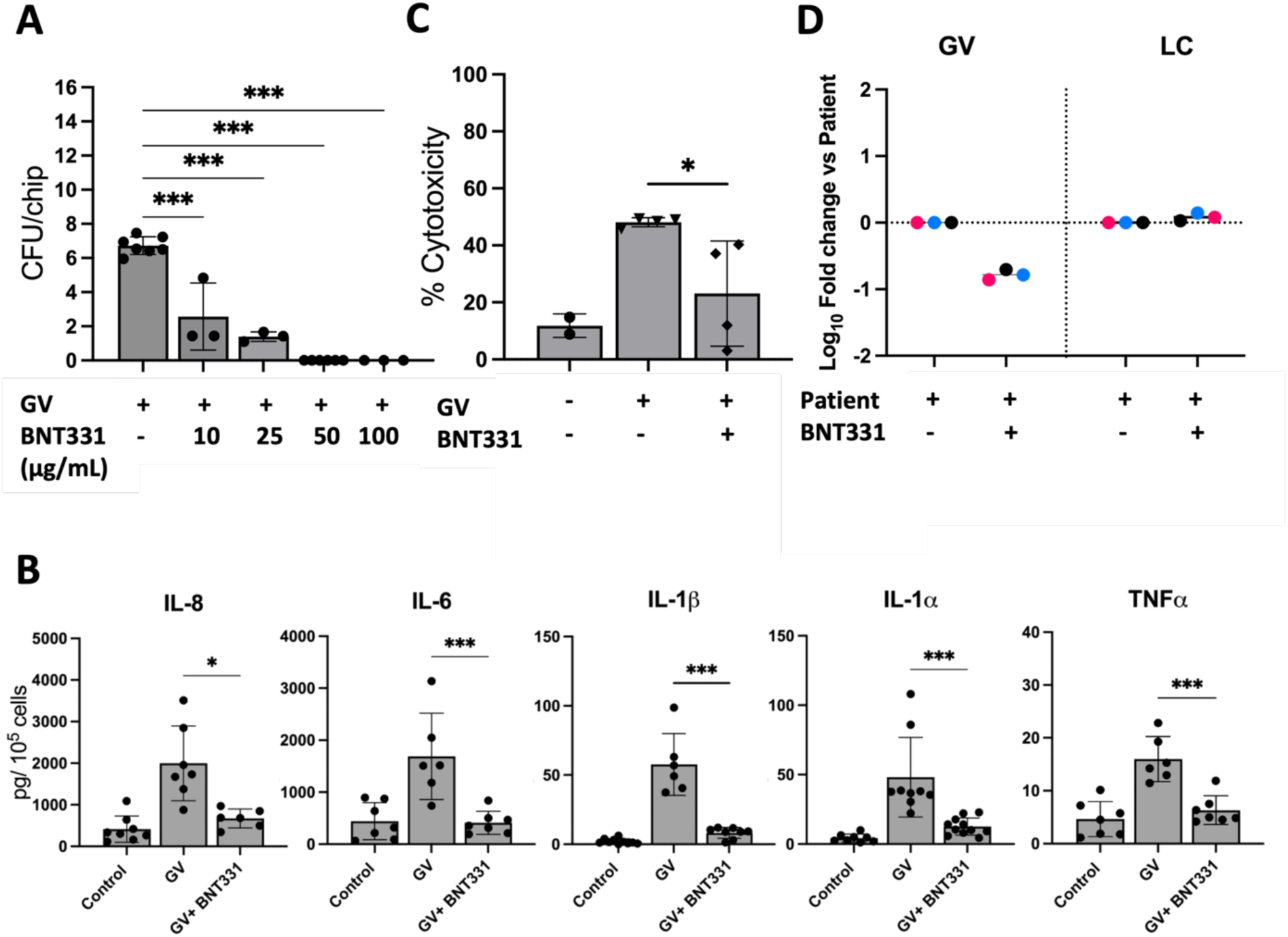
BNT331 kills *G. vaginalis* in the Vagina Chip. **A)** Vagina Chips inoculated with *G. vaginalis* (GV) were treated with different doses of BNT331 24 h after inoculation. CFU of GV was enumerated 72 h after BNT331 treatment. **B)** Pro-inflammatory cytokines were measured after 72 h in the outflows of Vagina Chips infected with GV and treated with BNT331 after 24 h of inoculation. Cytotoxicity was measured after 72 h in the outflows of Vagina Chips infected with GV and treated with 50 ug/mL BNT331 after 24h of inoculation. **(A-C)** Bar graphs represent Mean± SD for 3 independent experiments. P- values were calculated using one-way ANOVA. **D)** Vagina Chips were infected with vaginal swabs from three BV patients and treated with either BNT331 alone or in combination with LC. The relative abundance of GV and LC were measured at 72 h in the Chip digests by 16S rRNA sequencing and normalized to levels in the Vagina Chip infected with patient sample. Patient 1, 2 and 3 are depicted as 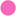, 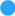 and 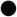 respectively.

Interestingly, the relative abundance of *L. crispatus* did not increase despite a 10-fold reduction in *G. vaginalis* upon BNT331 treatment (**Fig. 3D**).

### Enhanced efficacy of combined administration of L. crispatus and BNT331

Next, we combined both potential therapeutics and tested it in the Vagina Chip. To do this, we infected the Vagina Chips with a consortium of dysbiotic bacteria containing *G. vaginalis*, *Atopobium vaginae,* and *Prevotella bivia* (BVC1), which has been used previously to model BV in vitro (*18*). Before carrying out combination studies, we tested if BNT331 inhibits the engraftment of the *L. crispatus* consortia on-chip and confirmed that it does not (**fig. S1**). We then infected the Vagina Chips with BVC1 for 24 hours before exposing them to either BNT331, *L. crispatus* consortia, or a combination of both. We found that BNT331 alone and in combination with *L. crispatus* reduced the dysbiotic bacterial load in the chip compared to either control BVC1-infected Chips or chips treated with *L. crispatus* alone (**Fig. 4A**). Importantly, administration of both, in combination also significantly increased the engraftment of *L. crispatus* in the Vagina Chip relative to the treatment condition with *L. crispatus* alone (**Fig. 4B**). Thus, the combined treatment shifted the vaginal microbiome community to a *L. crispatus* dominant state unlike the chips treated with *L. crispatus* alone, which continued to exhibit a BVC1-dominant community (**Fig. 4C**). All treatments (BNT331, *L. crispatus* or both in combination) reduced BVC1-induced inflammation in the Vagina Chip (**Fig. 4D**). However, the increase in *L. crispatus* dominance induced by combined treatment with BNT331 and *L. crispatus* also resulted in an increase in D-lactate present in the Vagina Chip (**Fig. 4E**), indicating a shift towards a healthy vaginal environment. These results show that BNT331 can effectively kill *G. vaginalis* thereby aiding the engraftment of *L. crispatus* in the Vagina Chip. They also confirm that vaginal inflammation is sensitive to the presence of both *G. vaginalis* and *L. crispatus* in the Vagina Chip, and that their effects are separable.

**Fig. 4.**
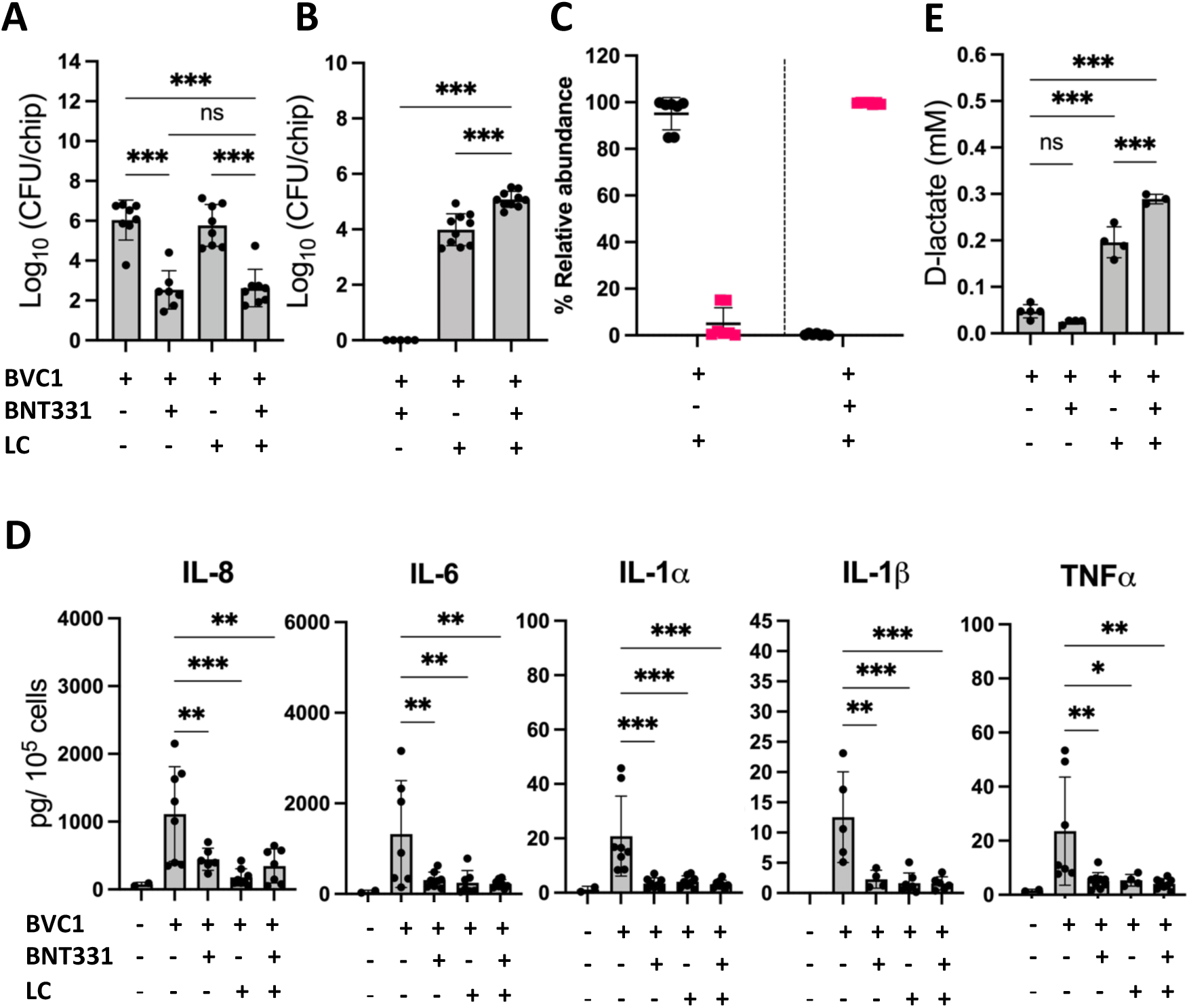
Combinational treatment of BNT331 and LC reduce dysbiosis in the Vagina Chip and helps restore a healthy vaginal state. A-B) Vagina Chips inoculated with BVC consortium and treated with 50ug/mL of BNT331 or LC or a combination of BNT331 and LC 24 h after inoculation. Bacterial load of BVC **(A**) and LC **(B)** was quantified by CFU enumeration 72 h after treatment. **C)** Relative abundance of BVC (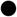) and LC (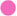) was calculated using CFU enumeration of bacteria in Vagina Chips infected with BVC for 24 h and then treated with either LC alone or in combination with BNT331 72 h after treatment. **D)** Pro-inflammatory cytokines measured in the outflows of Vagina Chips infected with BVC and treated with BNT331 or LC or a combination of BNT331 and LC after 72 h of treatment. **E)** D-lactate was measured in outflows from Vagina Chips inoculated with BVC consortium and treated with 50 ug/mL of BNT331 or LC or a combination of BNT331 and LC 72 h after treatment. **(A-B, D-E)** Bar graphs represent Mean± SD from three independent experiments. P-values were calculated using one-way ANOVA.

The vaginal microenvironment is complex and consists of a mix of diverse bacterial species. So, to probe whether a combination of BNT331 and *L. crispatus* would be effective in treating BV, we inoculated the Vagina Chips with swab samples containing a more complex vaginal microbiome from three BV patients for 1 day and then treated the chips with a combination of BNT331 and *L. crispatus*. After 72 h of treatment, the engrafted bacterial community in the Vagina Chip was analyzed by 16S rRNA sequencing (**fig. S2**). When the Vagina Chips infected with each of the three patient swabs were treated with a combination of BNT331 and *L. crispatus*, there was almost a 10-fold reduction in *G. vaginalis* and an increase in *L. crispatus* by 5 to 100- fold depending on the patient chip (**Fig. 5A**).The engraftment of *L. crispatus* induced by this therapeutic combination also was accompanied by an increase in D-lactate levels (**Fig. 5B**) and a reduction in inflammation (**Fig. 5C**) in the Vagina Chip, indicating restoration of a healthier vaginal microenvironment.

**Fig. 5.**
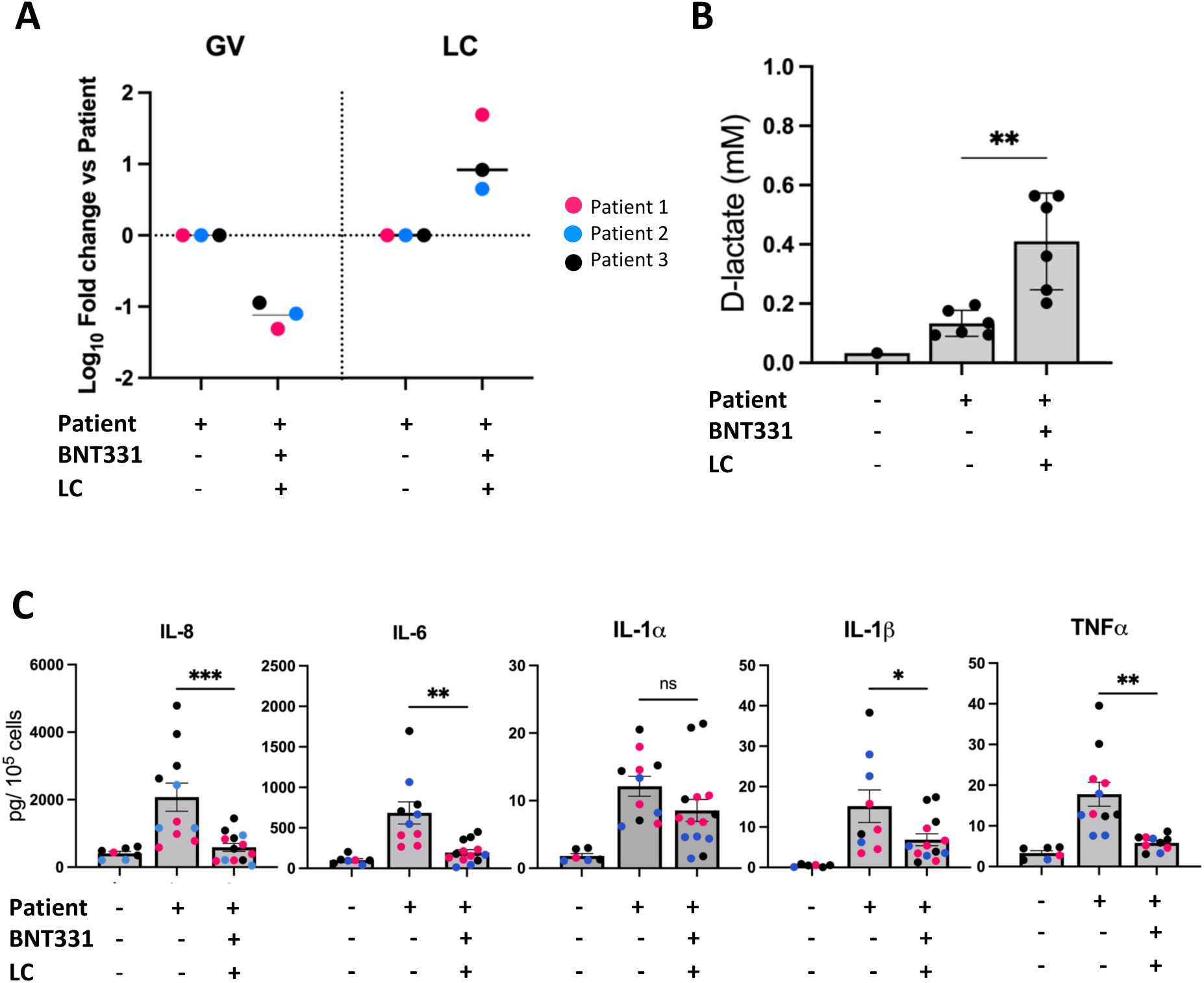
BNT331 reduces G. vaginalis and helps LC establish in Vagina Chips infected with vaginal swab samples from BV patients. **A)** Vagina Chips were infected with vaginal swabs from three patients with BV and treated with either BNT331 alone or in combination with LC. The relative abundance of GV and LC were measured by 16S rRNA sequencing and normalized to their levels in the Vagina Chip infected with Patient sample at 72 h. **B-C)** D-lactate **(B)** and pro-inflammatory cytokines **(C)** were analyzed from outflows from the Vagina Chips infected with vaginal swabs from three BV patients and treated with either BNT331 alone or in combination with LC. Bar graphs represent Mean± SD. P- values were calculated using one-way ANOVA. Patients 1, 2 and 3 are depicted as 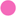, 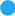 and 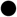 respectively.

## DISCUSSION

Bacterial vaginosis is a disease affecting about 30% women globally. The current standard of care treatment is based on administration of antibiotics, such as metronidazole or clindamycin. However, there is a great need for better BV therapeutics because many women experience disease recurrence within a year. Failure to fully treat BV is also associated with an increased risk of venereal diseases and preterm birth. In this study, we compared the therapeutic effects of a synthetic consortium of optimal strains of *L. crispatus* and the endolysin BNT331 using a human Vagina Chip BV model that contains a dysbiotic microbiome composed of either *G. vaginalis* alone or *G. vaginalis*, *A. vaginae,* and *P. bivia* (BVC1) in direct contact with the human vaginal epithelium (*18*). We found that the *L. crispatus* consortium reduced BV-induced inflammation in the Vagina Chip but did not reduce the bacterial load of the BVC1 consortium. When the Vagina Chips were infected with patient-derived consortia, we observed an overgrowth of several aerobic species such as *Escherichia, Streptococcus* and *Enterococcus* after 72h in all treatment conditions, which is atypical of BV. BNT331 effectively killed *G. vaginalis* in BV-infected Vagina Chips infected with the BVC1 consortium and the patient-derived consortia. It was not able to restore *L. crispatus* dominance in the patient-derived community, which may be related to the overgrowth of the aerobic species. Importantly, however, when we combined the BNT331 with *L. crispatus* consortium, we observed both a complete reduction in *G. vaginalis* and a concomitant increase in engraftment of *L. crispatus* bacteria. Taken together, these results show that when combined with a *L. crispatus* consortium, BNT331 may be more effective in shifting the bacterial community to a healthy state in the human Vagina Chip and potentially in women. These results suggest that a combination of BNT331 and an LBP formulation containing *L. crispatus sp.* may represent a more effective way to treat BV.

Characterization of the effects of culturing *G. vaginalis* and *L. crispatus* bacteria in the epithelial lumen of the human Vagina Chip confirmed that this model effectively recapitulates the dysbiotic environment produced by the former, as indicated by increased inflammation and cell injury as well as the health promoting effects of the latter, as shown by suppression of inflammation and cytotoxicity. Some groups are exploring the possibility of using a LBP composed of *L. crispatus* consortia for the treatment of BV based on the concept that shifting the vaginal microbiome to a *L. crispatus* dominated community could potentially reverse the disorder (*17, 20*). Our results show that while the administration of a *L. crispatus* consortium was effective at suppressing the inflammatory and cytotoxic effects of *G. vaginalis*, it did not effectively switch the microbiome into a healthy state as the *G. vaginalis* bacterial load remain high. This finding suggests that the addition of *L. crispatus* bacteria is able to alter the virulence of *G. vaginalis* without displacing it from the vaginal tissue. This is reminiscent of the past finding that a cell free supernatant of *L. plantarum* was able to alter the expression of virulence-related genes (e.g., vaginolysin) in *G. vaginalis* (*15*).

Our findings in the Vagina Chip suggest that while treatment with *L. crispatus* might benefit patients by reducing BV symptoms, it might not be an effective solution if the *L. crispatus* are not able to reestablish themselves as the dominant microbial community in the vaginal lumen. This is consistent with a past clinical study in which administration of a single strain of *L. crispatus* LBP (Lactin V) was complicated by recurrence of BV in ∼30 % of patients by week 12 (*17*). *G. vaginalis* forms biofilms on the vaginal epithelium (*21–23*) that are resistant to most antibiotics and their presence has been associated with an increased rate of recurrence of BV after metronidazole treatment (*24, 25*). Hence, another therapeutic approach for BV would be to develop drugs that kill *G. vaginalis* within biofilms so that it can be released from the vaginal epithelium. This is the rationale behind the development of the *G. vaginalis*-specific endolysin BNT331 that we tested in this study. Indeed, we found that BNT331 completely eradicated *G. vaginalis* from the epithelial lining of the Vagina Chip.

Consequently, it also reduced the inflammation and cell injury induced by *G. vaginalis*. To simulate more physiological conditions, we inoculated the Vagina Chips with vaginal swabs from BV patients. We found that BNT331 reduced *G. vaginalis* in these Vagina Chips, but this reduction did not result in restoration of a dominant *L. crispatus* community, probably due to low abundance of *L. crispatus* in the vaginal swab samples from BV patients used in this study and an overgrowth of fast-growing bacterial species (e.g., *Streptococcus*, *E. coli,* etc.) in the Vagina Chips, which is atypical of BV *in vivo*.

This is important because some reports have suggested that administration of *Lactobacillus* sp. reduces the recurrence rate of BV after treatment with metronidazole (*17, 20*).

While *G. vaginalis* is present in the vaginal microbiome of most women with BV, other anaerobic bacterial species are also often present. For this reason, we tested the effect of BNT331 in combination with *L. crispatus* in Vagina Chips infected with the synthetic dysbiotic consortium, BVC1, that contains *G. vaginalis*, *P. bivia* and *A. vaginae*, all of which are major players in BV. In addition, we carried out similar studies with complex bacterial consortia isolated from swab samples taken from BV patients. Interestingly, we found that BNT331 facilitated the engraftment of *L. crispatus* in BVC1- infected Vagina Chips and those infected with swab samples from BV patients thereby re-establishing a healthy vaginal microenvironment. More specifically, when we inoculated the luminal channel lined by vaginal epithelium in the Vagina Chips with swab samples from BV patients, we observed engraftment of most of the strains present in the swab sample, although there were some differences in the composition of the bacterial community that developed on-chip. This may be due to presence of fungal species such as *Candida albicans* and fast-growing bacterial species such as *Streptococcus* sp., *E. coli* etc. in the swabs collected from patients. Additionally, some reports have previously shown that *Candida* promotes the growth of *Streptococcus* (*26*) which may have also contributed to their overgrowth in the Vagina Chips. Future studies will be needed to eliminate fungal components and slow down the growth of fast-growing bacteria to better mimic the patient community on the Chip.

In summary, this study suggests that *L. crispatus* containing LBPs and therapies targeting a major BV causing bacterium, *G. vaginalis*, both may have significant therapeutic effects on the vaginal health under BV conditions. However, a combination of *L. crispatus* and BNT331 therapies may represent a more effective treatment for BV to both restore a healthy vaginal microenvironment and reduce recurrence. This study further highlights the potential of the Vagina Chip as a pre-clinical analysis tool to test therapeutics and other treatment strategies for diseases associated with vaginal dysbiosis.

## MATERIALS AND METHODS

### Vagina Chip Culture

Human Vagina Chips were created and cultured following a previously published protocol (*18, 27*). Briefly, the two-channel microfluidic organ chip devices were obtained from Emulate Inc. (Boston, MA). The chip devices were activated according to manufacturer’s protocol. Then, the apical channel of these organ chip devices was coated with a mixture of 200 ug/mL Collagen I (Corning, cat. No. 354236) and 30 ug/mL Collagen IV (Sigma, C7521) and the basal channel was coated with a mixture of 200 ug/mL Collagen I and 15 ug/mL Poly-L-Lysine (ScienCell Research Laboratories, cat no. 2301) and incubated at 37 °C for overnight. Then, the apical and basal coating was washed with 200 uL of Vaginal epithelial medium (Lifeline Cell technology, cat.no. LL- 0068) and Fibroblast medium (ScienCell Research Laboratories, cat. No. 2301) respectively. Then, the basal channel of the chips was seeded with 50 uL of 5 X10^5^ cells/mL of human uterine fibroblasts (ScienCell Research Laboratories, cat. no. 7040) and the chips were inverted and incubated for 1.5 hours at 37 °C, 5 % CO2. Then, 200 uL of vaginal epithelium growth medium and fibroblast growth medium were added respectively to apical and basal channels. The following day, the apical channel was seeded with 50 uL of 3 X 10^6^ cells/mL of primary vaginal epithelial cells (Lifeline Cell technology, cat. no. FC-0083) and incubated overnight at 37 °C, 5% CO2. The chips were then hooked to pods and were subjected to continuous flow at a flow rate of 30 uL/h in the basal channel and intermittent flow at a flow rate of 15 uL/h in the apical channel for 2 days. Then, the chips were subjected to differentiation for 8 days by flowing the apical channel with hank’s Balanced Salt Solution (HBSS; ThermoFischer,

Cat. No. 14025092) intermittently at 15 uL/h and basal channel with in-house prepared differentiation media (4 mM L-glutamine, 20 mM Hydrocortisone, 1X ITES, 20 nM Triiodothyronine, 100 μM O-Phosphoryl Ethanolamine, 180 μM Adenine, 3.2 mM Calcium Chloride, 2 % Heat-inactivated FBS, 1 % Pen-strep, 4 nM Estradiol and 24 % Ham’s F-12 media to Low glucose DMEM) and flowing continuously at 30 uL/h. The apical and basal media of the Vagina Chips were changed to HBSS (LB/+G) (1.26 mM Calcium chloride, 0.49 mM Magnesium chloride hexahydrate, 0.406 mM Magnesium sulphate, 5.33 mM Potassium chloride, 137.93 mM Sodium chloride, 0.441 mM Potassium phosphate monobasic, and 5.55 mM D- Glucose to dd H2O; pH 4.8) and PS free differentiation media respectively 24 hours before introduction of bacteria.

### Lactobacillus crispatus consortium studies

To mimic a healthy vaginal microenvironment, three clinical isolates of *L. crispatus* namely C0059A1, C0124A1 and C0175A1 obtained from women participants of the UMB-HMP study with a stable *L. crispatus* dominant vaginal microbiome were mixed together to form a consortium. Each of the strains were grown overnight in Man, Rogosa, and Sharpe (MRS) broth (Fisher Scientific, cat. No. 288210) at 37 °C in an anaerobic chamber under 83% N2, 10 % CO2, 7% H2. To make stocks for inoculation, the culture was aliquoted, mixed with sterile glycerol (MP Biomedicals, Cat. No. 76019- 966) at a concentration of 16 % and frozen at -80 °C. A single aliquot was plated on MRS agar (Hardy, Cat no. G117) and incubated for 48 h at 37 °C under anaerobic conditions to calculate the colony forming units (CFU)/mL for each strain individually.

To form the consortium, required volumes of C0059A1, C00124A1 and C00175A1 stocks were mixed to achieve equal bacterial numbers per mL of inoculum at a final density of ∼ 10^6^/ mL. The bacterial mix was then washed and resuspended in HBSS (LB/+G) to prepare the inoculum.

### Dysbiotic *G. vaginalis* and BVC1 consortia studies

Clinical isolates of *G. vaginalis* (C0011E2 and C0011E4), *P. bivia* (0795_578_1_1_BHK8) and *A. vaginae* (0795_578_1_1_BHK4) were obtained from women with BV who participated in UMB-HMP (*28*) and Females Rising Through Education Support and Health study (*29*). Each of these strains was grown in peptone, yeast and tryptone (with Hemin and Vitamin K1) broth at 37 °C under anerobic conditions (83% N2, 10 % CO2, 7% H2). To make stocks for inoculation, the culture was aliquoted, mixed with sterile glycerol (MP Biomedicals, Cat. No. 76019-966) at a concentration of 16 % and frozen at -80 C. A single aliquot was plated on Brucella blood agar (with Hemin and vitamin K1) (Hardy, Cat no. W23) and incubated for 48 h at 37 °C under anaerobic conditions to calculate the colony forming units (CFU)/mL for each strain individually.

In this study we used either a consortium of two *G. vaginalis* strains (C0011E2 and C0011E4 or the previously published BVC1 consortium (*18*) containing *G. vaginalis* (C0011E2 and C0011E4), *P. bivia* (0795_578_1_1_BHK8) and *A. vaginae* (0795_578_1_1_BHK4). To form each of the consortium, required volumes of each stock was mixed to achieve equal bacterial numbers per mL of inoculum at a final density of ∼ 10^6^/mL. The bacterial mix was then washed and resuspended in HBSS (LB/+G) to prepare the inoculum.

### Co-culture of Vagina Chips with living bacteria

The apical and basal media of the Vagina Chips were changed to HBSS (LB/+G) and PS free differentiation medium, respectively, 24 hours before introduction of bacteria. For the infection with synthetic bacterial consortia, 37 uL of the inoculum was added to the apical channel of the Vagina Chips and the chips were incubated statically at 37 °C, 5 % CO2 for 20 h. Then, the chips were subjected to a flow rate of 40 uL/h with intermittent flow for 4 h per day in the apical channel and continuous flow in the basal channel. The outflows were collected at 24 h, 48 h and 72 h of infection and used for analysis. Vagina Chips were digested at 72 h using 1 mg/mL collagenase IV (Gibco, cat no. 17104019) in TrypLE (ThermoFisher, cat no. 12605010) to assess the number of adherent bacteria in the chips.

For infection with patient-derived vaginal microbiome samples, vaginal swabs from patients with BV were collected as part of a study at Massachusetts General Hospital (Partners Institutional Review Board, IRB number-2020P00393) (*30*). At each visit, participants self-collected vaginal swabs by inserting the swab 4-5cm into the vagina, rotating several times and then removing and placing in a sterile cryovial containing 10% glycerol/thioglycolate solution. The swab shaft was cut off with sterile scissors and then the sample was transferred to -80 C until thawed for use. For the inoculum, 250 uL of the swab was washed and resuspended in HBSS (LB/+G) and kept on ice. Then, 37 uL of the inoculum was added to the apical channel of the Vagina

Chips and the chips were incubated statically at 37 °C, 5 % CO2 for 20 h. Then, the chips were subjected to a flow rate of 40 uL/h with intermittent flow for 4 h per day in the apical channel and continuous flow in the basal channel. The outflows were collected at 24 h, 48 h and 72 h of infection and used for analysis. Vagina Chips were digested at 72 h using 1 mg/mL collagenase IV (Gibco, cat no. 17104019) in TrypLE (ThermoFisher, cat no. 12605010) to assess the adherent bacteria in the chips.

### Live Biotherapeutics (LBP) studies

For studies with *Lactobacillus crispatus* containing LBP consortium containing three clinical isolates of *L. crispatus* namely C0059A1, C0124A1 and C0175A1, the bacterial inoculum at a final density of ∼ 10^6^/ mL was prepared as described above. 37 uL of *L. crispatus* inoculum was added apically to Vagina Chips 24 h after inoculation with either G. vaginalis or BVC1 consortium and 37 uL of HBSS (LB/+G) was added apically to all control chips. The chips were then incubated statically for 20 h before being hooked to flow with HBSS (LB/+G, but without *L. crispatus*) in apical channel and PS free differentiation medium in basal channels at a flow rate of 40 uL/h. The apical channel was flown intermittently for 4 h each day and the basal channel was flown continuously. The outflows were collected each day after the intermittent flow until 72 h.

### BNT331 Studies

BNT331 was reconstituted in HBSS (LB/+G) at a concentration of 1 mg/mL and filtered through a 0.22 um syringe filter. It was then diluted in HBSS (LB/+G) to the desired concentration before being added to the chips. 37 uL of BNT331 at the desired concentration was added apically to the Vagina Chips 24 h after inoculation with *L. crispatus*, *G. vaginalis* or BVC consortia. In cases of a combination treatment with BNT331 and *L. crispatus* consortium, *L. crispatus* was resuspended in HBSS (LB/+G) containing BNT331. The chips were incubated statically for 20 h before being hooked to flow with HBSS (LB/+G) containing BNT331 in apical channel and PS free differentiation medium in basal channels at a flow rate of 40 uL/h. The apical channel was flown intermittently for 4 h each day and the basal channel was flown continuously. The outflows were collected each day after the intermittent flow until 72 h.

### Quantification of bacterial load

Effluents (50 uL) from the outlet of the epithelial channel collected at 24, 48 and 72 h and chip digests collected at 72 h were stored at -80 C with 16 % Glycerol. To enumerate the *L. crispatus* in the Vagina Chips, the effluents and digests were plated on MRS agar and incubated at 37 °C for 48 h under anerobic conditions. The colonies thus obtained in both effluents and chip digests were counted and total CFU was enumerated for each chip. To enumerate the non-optimal bacteria in the Vagina Chips, the effluents and digests were plated on Brucella blood agar (with Hemin and vitamin K1) (BBA) and incubated at 37 °C for 48 h under anaerobic conditions. The colonies thus obtained in both effluents and chip digests were counted and total CFU/chip was calculated.

### Immunofluorescence microscopy

Bacteria were stained with Bactoview (Biotium, cat. No. 40101) according to manufacturer’s protocol and washed with HBSS (LB/+G) twice before inoculation of the Vagina Chips. After 72 h of inoculation, Vagina Chips were fixed with 4 % paraformaldehyde (Alfa Aesar, cat. No. J61899) at room temperature for 1 h and then the chips were washed with DPBS -/- (cat. No. 14190-144). The chips were then permeabilized using 0.1 % Triton-X-100 (Sigma-Aldrich, cat. no. 9036-19-5) in DPBS -/-, blocked for 1 h with 5 % goat serum at room temperature and then, incubated with primary antibody against ZO-1 (Abcam, cat. No. ab276131 at 1:100 dilution) for overnight at 4 C. Then, the chips were washed thrice with DPBS -/- and incubated with secondary antibody (Abcam, cat. No. ab150077 at a dilution of 1:500) and Alexa-fluor tagged antibody against CK-5 (Abcam, cat. No. ab193894 at a dilution of 1:200) for 2 h at room temperature in the dark. Then the chips were washed twice with DPBS -/- and counterstained with DAPI (Invitrogen, cat. No. D1306) at a concentration of 1 ug/mL for 5 mins at room temperature. Images were acquired with an inverted laser-scanning confocal microscope (Leica SP5 X MP DMI-6000) and processed using Image J/Fiji.

### Cytokine and chemokine analysis

The apical effluents collected from Vagina chips were analyzed for IL-8, IL-6, TNF-α, IL-1α, IL-1ϕ3, IFN-g, IL-10, MIP-1α, MIP-1ϕ3, IP-10 and RANTES using customized ProcartaPlex assay kits (ThermoFisher Scientific) and the concentration of the analytes was measured using a Luminex 100/200 Flexmap 3D instrument and analyzed using the Luminex XPONENT software.

### Cytotoxicity analysis

The apical effluents collected from Vagina chips at 72 h were analyzed for cytotoxicity using the LDH assay kit (Promega, cat. no. G1780) according to manufacturer’s protocol. For the positive control, the chip digest of an uninfected control chip was lysed using 10X lysis buffer provided with the kit.

### D-Lactate analysis

The apical effluents collected from the Vagina Chips were collected and equilibrated in an anaerobic chamber under 83% N2, 10 % CO2, 7% H2. D-lactate in the effluents was measured using EnzyChrom D-Lactate assay kit (BioAssay Systems, cat. No. EDLC-100) according to manufacturer’s protocol.

### 16S rRNA gene-based microbial community analysis

To assess bacteria adhered to the vaginal epithelial cells, total nucleic acids (TNA) were extracted from vagina chip digests using a phenol-chloroform method, which includes a bead beating process to disrupt bacteria (*29, 31*). Samples were transferred into a solution containing phenol: chloroform: isoamyl alcohol (PCI, 25:24:1, pH 7.0) and 20% sodium dodecyl sulfate in Tris-EDTA buffer with sterile 0.1mm glass beads and homogenized on a bead beater for 2 min at 4C. After centrifuged at 6800 xg for 3 min at 4 C, the top aqueous phase was transferred to a clean tube with equal volume of PCI, mixed and centrifuged at 16000 xg for 5 min at 4C. The aqueous phase was transferred to a clean tube and precipitated with isopropanol and sodium acetate overnight at -20 C. Samples were then centrifuged at 21000 xg for 30 min at 4C, supernatant was removed, and pellet was washed with 100% ethanol. After centrifuge at 21000 xg for 15min at 4C, the ethanol was discarded and the tube was left open to dry at room temperature. TNA sample was resuspended in 20uL of Tris-EDTA buffer.

Bacterial taxonomic compositions in samples were determined through sequencing the 16S rRNA gene V4 region. This region was amplified using the primer set 515F/806R (with 806R barcoded for multiplexing) in a 25 ul reaction containing 1X Q5 reaction buffer (NEB), 0.2 mM of dNTPs, 0.2 uM of each primer, 0.5 unit of Q5 high-fidelity DNA polymerase (NEB), and 2 ul of TNA sample. PCR was performed in triplicate for each sample at 98°C for 30 s, followed by30 cycles of 98°C for 10s, 60°C for 30s, and 72°C for 20s, with a final extension at 72°C for 2 min. To monitor contamination, negative controls with water as template, in parallel to TNA samples in each barcode master mix, were also PCR amplified to confirm no amplification. Triplicate PCRs for each sample were combined and checked on an agarose gel. PCR products of samples were pooled based on gel band strength, purified with QIAquick PCR purification kit, and sequenced on Illumina MiSeq with a 300-cycle kit. Negative controls for TNA extraction and PCR were included in the sequencing.

Denoising and removal of sequencing errors from the Illumina amplicon reads was conducted using DADA2 (version 1.26). Taxonomic assignments were made with Genome Taxonomy Database (version R207) and curated taxonomy from prior vaginal microbiome studies (*29*). The data are represented as relative abundance of *G. vaginalis* and *L. crispatus* in Vagina Chips treated with BNT331 alone or in combination with *L. crispatus* normalized to the relative abundance in patient swab-infected Vagina Chips.

### Statistical analysis

All results were obtained from at least three independent experiments and are represented as mean ± standard deviation (SD). Statistical significance was calculated using one-way ANOVA using GraphPad Prism.

## Acknowledgments

We thank Jacques Ravel, Ph.D, and Seth Rakoff-Nahoum, Ph.D., for providing the bacterial strains of *L. crispatus, G. vaginalis, P. bivia* and *A. vaginae* used in this study.

## Funding

This research was sponsored by funding from the Bill and Melinda Gates Foundation (INV-035977) and the Wyss Institute for Biologically Inspired Engineering at Harvard University.

## Author contributions

Conceptualization: AG, DEI; Methodology development: AG, AJ, LPT, LC; Investigation and data analysis: AG, ACJ, AW, HR, MFZ, JX, JE, OG, AS, NL, YM; Patient swab collection: BC, CM; Visualization: AG, AW; Funding acquisition: AG, AJ, GG, DEI; Project administration: AJ; Supervision: DEI, DK, CM; Writing – original draft: AG, DEI; Writing – review & editing: All authors.

## Competing interests

D.E.I. holds equity in Emulate, chairs its scientific advisory board and is a member of its board of directors. L.C. and L.P.T. are employees of BioNTech and inventors on several patent applications related to BNT331. CM has a financial interest in Ancilia Biosciences, a company developing a new class of Live Biotherapeutics and other bacterial products. CM’s interests were reviewed and are managed by MGH and Mass General Brigham in accordance with their conflict-of-interest policies. CM is on the Scientific Advisory Board of Concerto Biosciences and has served as a consultant for Freya Biosciences. The remaining authors declare no competing interests.

## Data and materials availability

All data are available in the main text or the supplementary materials.

## Declarations

This manuscript does not contain any individual person’s data in any form. Hence, the consent for publication is not applicable to this study.

## Supplementary Materials

**Fig. S1.**
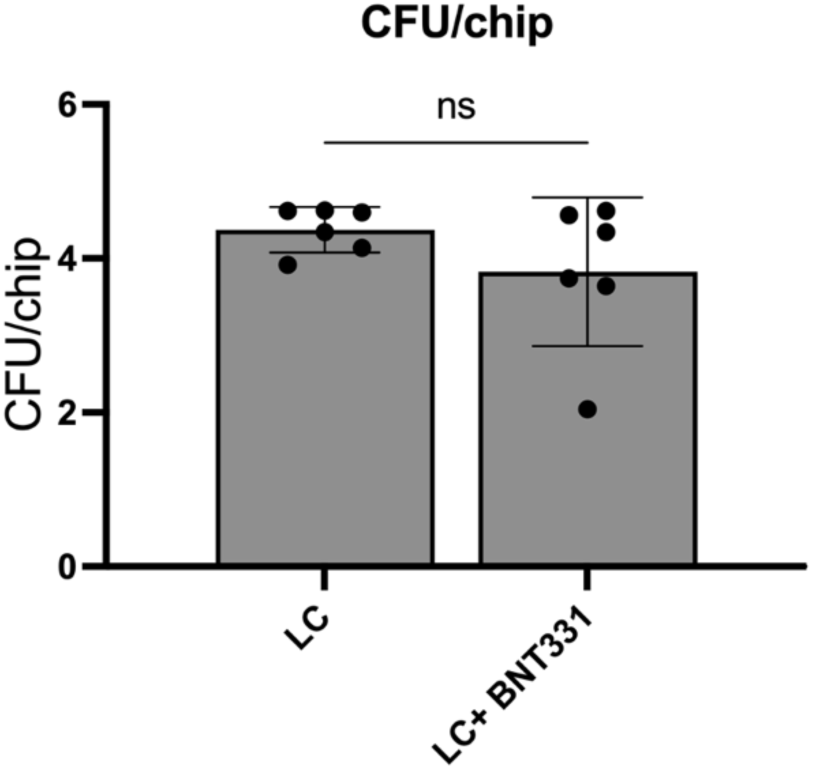
Effect of BNT331 on engraftment of *L. crispatus* in Vagina Chips. Vagina Chips inoculated with *L. crispatus* consortium (LC) were treated with 50 ug/mL of BNT331 24 h after inoculation. CFU of LC was enumerated 72 h after BNT331 treatment.

**Fig. S2.**
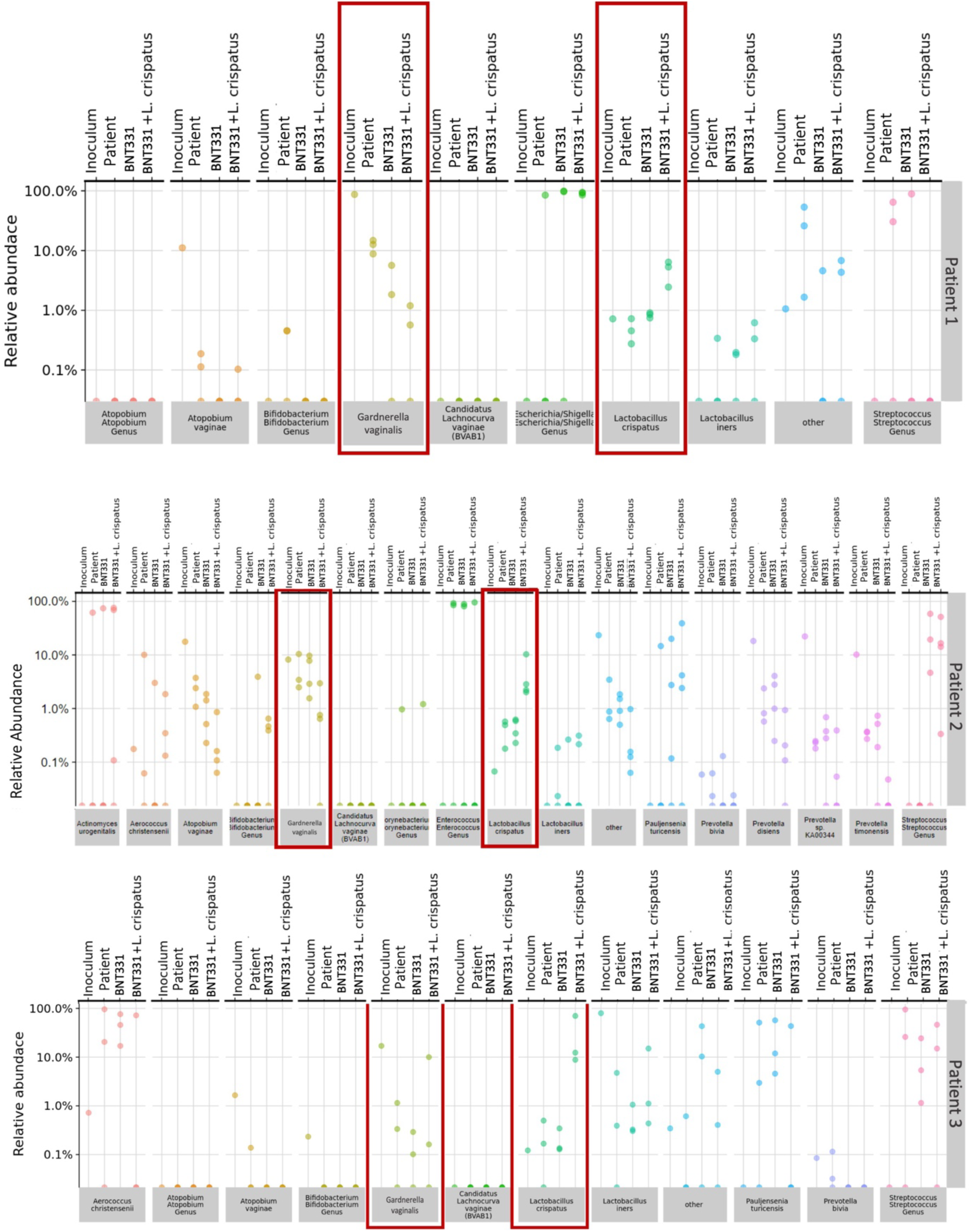
16S rRNA sequencing of bacteria in the Vagina Chips infected with patient samples upon treatment with BNT331 and LC. Vagina Chips inoculated with swabs from three BV patients were treated with BNT331 alone or in combination with *L. crispatus* containing LBP 24 h after inoculation. The relative abundance of bacterial species engrafted in these infected Vagina Chips after 72 h of treatment were detected using 16S rRNA sequencing. Inoculum represents the bacterial community in the patient swab before adding to the Vagina Chip; Patient represents the bacterial community that developed on chip after 72 h of buffer treatment.

